# GI-NemaTracker - A farm system-level mathematical model to predict the consequences of gastrointestinal parasite control strategies in sheep

**DOI:** 10.1101/2025.04.01.646550

**Authors:** Lee Benson, Ilias Kyriazakis, Naomi Fox, Alison Howell, Giles T Innocent, Fiona Kenyon, Diana Williams, David A Ewing

## Abstract

Gastro-intestinal nematode infections are considered one of the major endemic diseases of sheep on the grounds of animal health and economic burden, both in the British Isles and globally. Parasites are increasingly developing resistance to commonly used anthelmintic treatments meaning that alternative control strategies that reduce or replace the use of anthelmintics are required. We present GI-NemaTracker, a systems-level mathematical model of the full host-parasite-environment system governing gastro-intestinal nematode transmission on a sheep farm. The model is based on a series of time-varying delay-differential equations that explicitly capture environmentally-driven time delays in nematode development. By taking a farm systems-level approach we represent both in-host and environmentally-driven free-living parasite dynamics and their interaction with a population of individually modelled lambs with diverse trait parameters assigned at birth. Thus we capture seasonally varying rates of parasite transmission and consequently variable weight gain of individual lambs throughout the season. The model is parameterised for *Teladorsagia circumcincta*, although the framework described could be applied to a range of nematode parasite species. We validate the model against experimental and field data and apply it to study the efficacy of four different anthelmintic treatment regimes (neo-suppresive treatment, strategic prophylactic treatment, treatment based on faecal egg counts and a regime which leaves 10% of the animals untreated) on lamb weight gain and pasture contamination. The model predicts that similar body weights at a flock level can be achieved while reducing the number of treatments administered, thus supporting a health plan that reduces anthelmintic treatments. As the model is capable of combining parasitic and free-living stages of the parasite with host performance, it is well suited to predict complex system responses under non-stationary conditions. The implications of the model and its potential as a tool in the development of sustainable control strategies in sheep are discussed.

## 1. Introduction

Parasitic diseases are ubiquitous and compromise the health, productivity and welfare of farmed animals. Gastro-intestinal nematode (GIN) infections can result in affected animals exhibiting a reduction in feed intake, redirection of nutritional resources and consequent reductions in weight gain of approximately 20% with high parasite burdens resulting in clinical gastro-enteritis (Mavrot et al., 2015). Beyond the obvious negative impact on animal health and welfare this threatens the economic viability of livestock production with estimated economic costs of € 1.8 billion per year across 18 European countries, with € 357 million of these losses occuring in sheep (Charlier et al., 2020). Infection has also been shown to increase the greenhouse gas emissions of affected sheep, with GIN infection estimated to increase peak methane yield in parasitised lambs by 33% relative to unparasitised lambs (Fox et al., 2018). Further, the ongoing effects of climate change are anticipated to increase the burden which GIN populations place on livestock due to increased temperatures and changing rainfall patterns creating favourable conditions for parasite development (Rose et al., 2015; Fox et al., 2015). Anthelmintics remain widely used to control GIN populations in livestock; however, parasite populations are increasingly developing resistance to anthelmintic treatments (Learmount et al., 2016) anthelmintics themselves have negative environmental impacts and harm non-target organisms (Vokřál et al., 2023). As such, we must seek novel methods by which the use of anthelmintics can be limited whilst the burden of GIN infection on livestock populations is reduced.

The principal driver to reduce the quantities of anthelmintics used is generally considered to be the emergence of anthelmintic resistance (AR), which was recently estimated to cost sheep farming € 10.5 million per year across 18 European countries due to lost productivity and the cost of partially ineffective anthelmintic drugs (Charlier et al., 2020). This cost is likely to increase given that AR has been increasing in the UK since 1980, with a recent study estimating resistance to four main classes of anthelmintic lies between 20 and 60% in sheep (Rose Vineer et al., 2020). In addition, there is growing evidence that anthelmintics can have a serious impact on the environment and cause harm to a wide range of invertebrate species, including dung fauna, causing damage to soil health (Sutton et al., 2014; Cooke et al., 2017; Vokřál et al., 2023). The extent of anthelmintic contamination is known to be widespread and to vary substantially by region (Vokřál et al., 2023) but the ecotoxological effects of are not yet fully understood (Mooney et al., 2021).

For some years industry bodies have been shifting their messaging away from anthelmintic dosing at set intervals to minimise parasite burdens, moving towards the idea of treating strategically to maintain a parasite population that is susceptible to anthelmintics without detriment to health and productivity (Mitchell, 2005; Stubbings et al., 2020). Reduction in the use of anthelmintics to manage GIN infection can be achieved through developing integrated control strategies based on improved diagnosis and farm management options (Mitchell, 2005) or targeting treatment at infected animals to treat disease or to stop super shedders spreading infection (Kenyon et al., 2013). Studies that have fully evaluated selective treatment programmes in sheep are limited and their findings are specific to the circumstances experienced in those studies (Cringoli et al., 2009; Kenyon and Jackson, 2012; Kenyon et al., 2013; Busin et al., 2013; Calvete et al., 2020). Given the notable diversity of farming systems and extensive spatio-temporal variation in climate, developing evidence-based, bespoke advice is difficult. Mathematical models are a valuable tool by which a range of novel control strategies can be explored before expensive farm trials (Rose Vineer et al., 2020).

Here we build on decades of empirical research and mathematical modelling on component parts of this system (Vagenas et al., 2007a; Laurenson et al., 2011; Rose et al., 2015; Rose Vineer et al., 2020; Filipe et al., 2023) to develop a systems model capable of addressing pressing questions related to control of GIN and development of AR. We develop a mathematical model which captures parasite dynamics both in host and on pasture and predicts the parasite burdens of individual lambs. Our model is based on delay-differential equations (DDEs) which capture the environmentally-driven time delays in development of the stage-structured nematode population more accurately than commonly used ordinary differential equations (ODEs). By modelling the interaction between a phenotypically diverse population of lambs and the environmentally-driven parasite populations we can predict body weight gain of individual lambs over the course of their first grazing season. We show that our systems-level approach produces outputs consistent with empirical evidence. It also enables, for the first time, in silico assessment of a range of strategies that were co-constructed with farmers over a series of focus groups to better manage GIN infections and AR in the real world. Our results show that strategies which reduce anthelmintic treatments by targeting individual animals or treating at particular times can achieve weight gains at a flock level that are comparable with repeated, calendar-based treatments throughout the season and point toward an opportunity to reduce the development of anthelmintic resistance.

## 2. Materials and methods

### 2.1. Model overview

We develop GI-NemaTracker, a mathematical model based principally on delay differential equations (DDEs) to model GIN transmission on a sheep farm. We explicitly model parasite, host and environment using linked equations, facilitating consideration of a variety of different anthelmintic treatment regimes that may target individual animals or groups of animals to allow exploration of the impacts of novel control scenarios. We parameterise our model based on *Teladorsagia circumcincta* and texel cross sheep in a typical lowland farm environment.

The model represents the full life cycle as GINs progress from the egg stage, *E*(*t*), through the pre-infective larvae stage, *L*_12_(*t*), to become infective larvae in faeces, *L*_3*f*_ (*t*), then infective larvae on pasture, *L*_3*p*_(*t*), which are further split between those on the soil, *L*_3*s*_(*t*) and those on herbage, *L*_3*h*_(*t*) (Figure 1).

**Figure 1:**
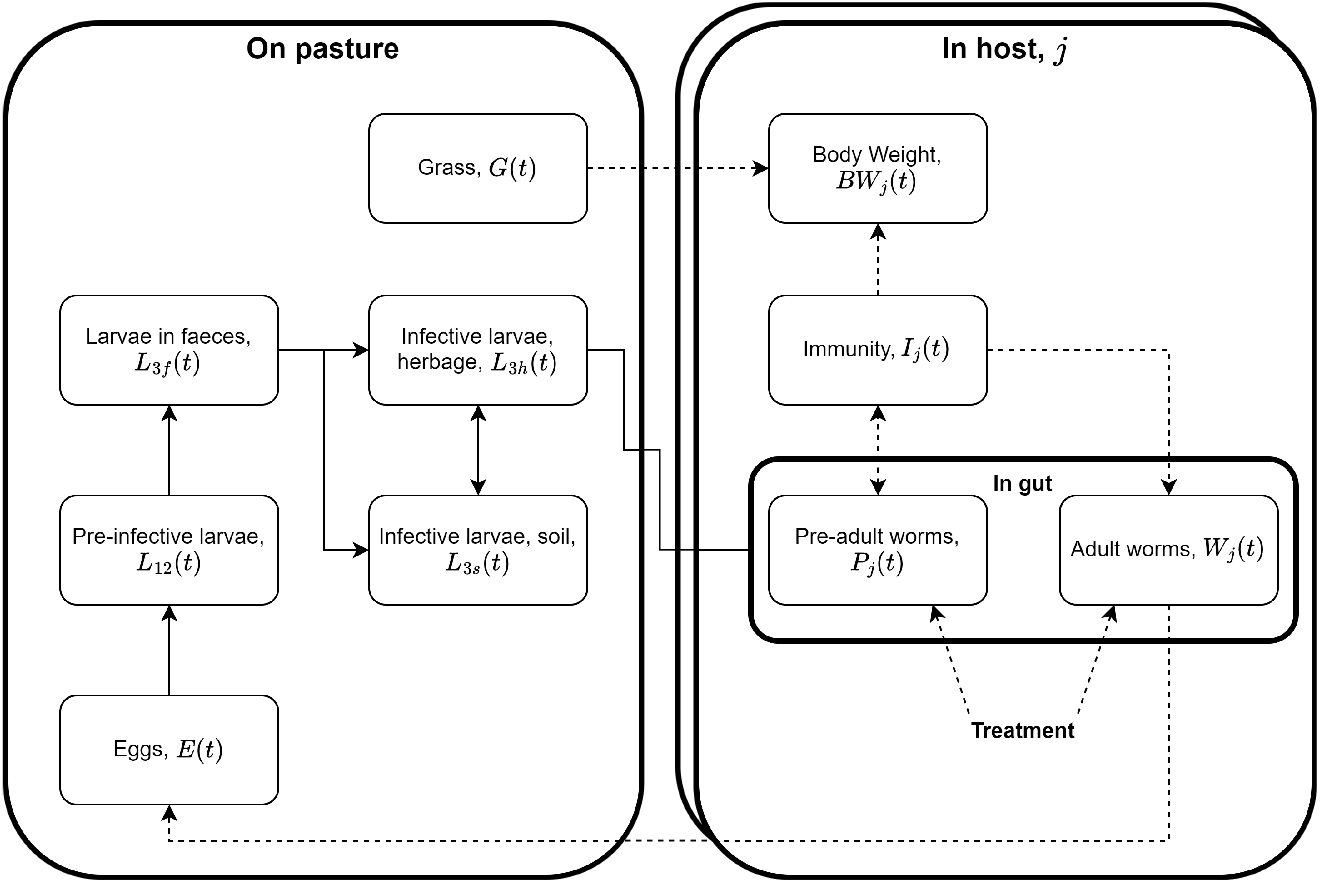
Model flowchart: The solid lines show direct progressions through the nematode life stages. The dashed lines show more indirect links between the model compartments. For example, solid lines between eggs, *E*(*t*) and pre-infective larvae, *L*_12_(*t*) represents the hatching of larvae from eggs, whilst the dashed line between infective larvae on herbage and immunity represents the effect of larval ingestion on the immunity level. The model representation shows a hypothetical animal, *j* and its interaction with nematodes on pasture, though multiple host animals (represented by the duplication of the rectangle) will all interact with the pasture simultaneously.

**Figure 2:**
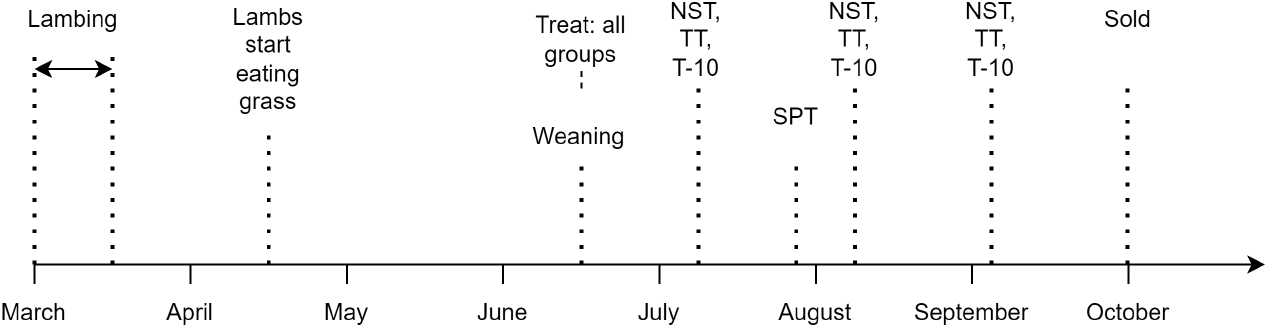
Timeline of events: the figure shows the assumed timeline of events under which we evaluate the 4 described anthelmintic treatment regimes and represents a group of 60 lambs with their mothers throughout the grazing season at a density of 24 lambs per hectare. The mothers are assumed not to contribute to parasite transmission due to immunity gained in previous years. The lines indicate treatment for the NST, SPT and TT-10 scenarios. For the TT scenario, it indicates a FEC and possible treatment if the threshold was reached.

The rate of progression and survival through these stages is dependent on environmental conditions. Infective larvae on herbage are then ingested by grazing lambs where they become pre-adult worms, *P*_*j*_(*t*), and some time later progress to the adult worm stage, *W*_*j*_(*t*), in host *j*. The establishment of pre-adults and the mortality and fecundity of adult worms are all affected by the immunity level of the host, *I*_*j*_(*t*), which develops in response to the ingestion of infectious larvae. Hosts will gain or lose body weight, *BW*_*j*_(*t*) at a rate which is determined by their maturity, their state of immunity (nutrient resource costs of gaining and maintaining immunity reduce the resources available for growth) and any parasite-induced reduction in feed intake (anorexia) exhibited as a result of nematode infestation. Treatment with anthelmintics acts as a short pulse which removes both the pre-adult and adult life stages. We assume that ewes have a negligible impact on the parasite burden on pasture due to immunity acquired in previous years so we only consider the worm burden in lambs from weaning onwards.

The model presented is principally based on the use of DDEs. Ordinary differential equations (ODEs) are very widely used and can be relatively straightforward to solve but they assume an exponential distribution of waiting times in each model compartment, which gives a poor reflection of the true duration of most invertebrate life stages and can consequently introduce bias in model estimates (Wearing et al., 2005). The linear chain trick, employed by Filipe et al. (2023) to model the pre-adult stage duration within the gut and the delay in the onset of immunity, allows a more realistic representation of the stage durations by linking model compartments so that the duration of each life stage is given by an Erlang distribution where the shape parameter is given by the number of linked compartments (Hurtado and Kirosingh, 2019). This gives a more realistic representation of the duration spent within each life stage; however, as model complexity increases the number of sub-compartments and thus the number of equations becomes unwieldy. This issue of increasing model complexity (and consequently increasing computational demands) is particularly pertinent given our aim to explicitly model individual hosts. The inclusion of mortality in any of the model life stages can also skew the distribution of waiting times when using the linear chain trick as individuals which would have been longer-lived are impacted more greatly by the mortality rate (Vansickle, 1977).

DDEs have been widely used to model insect populations using the framework first popularised by Gurney et al. (1983) and Nisbet and Gurney (1983) and provide a compromise whereby all individuals which enter a life stage together at time *t* will leave the stage together at time *t* + τ where τ is the duration of the life stage. This assumption is perhaps less realistic than the Erlang distribution generated by the linear chain trick but it is much more tractable and allows us to feasibly model the parasite burden of each individual sheep and each patch of pasture on the farm whilst still explicitly modelling the delays in nematode development both in host and on pasture.

### 2.2. Full model formulation

In this section we present a full description of the model formulation based on coupled-delay differential equations representing the transmission of GINs between a diverse host population and the environment. In the model formulation below we do not explicitly represent which of the model parameters are considered to vary between hosts and which are held constant for all hosts because in principle any of the parameters could be allowed to vary between hosts. Those parameters which we have assumed to be constant are given in Table 1, those assumed to show inter-host variability are given in Table 2 and the functional forms describing the environmental impacts on free-living nematodes are given in Table 3.

**Table 1:** Parameter values that were held constant across all lambs with a description of the parameter and the reference.

| Parameter | Description | Value | Reference<sup>a</sup> |
| --- | --- | --- | --- |
| $BW_m$ | mature bodyweight | 55kg | Assumed<sup>b</sup> |
| $d$ | digestibility of dry matter | 0.756 | [1] |
| $a_w$ | allometric constant | $1.377 \text{ kg}^{1-b_w}$ | [2] |
| $b_w$ | allometric constant | 0.741 | [2] |
| $p_e$ | prop. of BW, excluding gut contents | 0.91 | [3] |
| $c_{I1}$ | acquisition of immunity | 25.0 | [4] |
| $\delta_{P0}, \delta_{P1}$ | pre-adult mortality rates at $I = 0, 1$ | 0.0152 per day, 0.1563 per day | [5] |
| $\delta_{W0}, \delta_{W1}$ | adult worm mortality rates at $I = 0, 1$ | 0.01 per day, 0.12 per day | [5] |
| $\delta_I$ | rate of waning of immunity | 0.002 per day | [6] |
| $\tau_I$ | delay in immune response | 7 days | [7] |
| $\tau_P$ | stage duration of in-host pre-adult larvae | 18.0 days | [8] |
| $p_f$ | female proportion of adult worms | 0.5 | [6] |
| $\Lambda$ | scale parameter FI reduction | 12 | Assumed |
| pFaecDM | DM proportion of faeces | 0.3 | [9] |
| $K_g$ | DM carry capacity of pasture | 3500 kg per hectare | [4] |
| $r_g$ | pasture DM growth rate | 70 kg per hectare per day | [4] |

<sup>a</sup> [1] - Martins et al. (2019), [2] - Galvani et al. (2008), [3] - Li et al. (2021), [4] - Filipe et al. (2023), [5] - Laurenson et al. (2011), [6] - Rose Vineer et al. (2020), [7] - Singleton et al. (2011), [8] - Stear et al. (1995), [9] - Le Jambre et al. (2007)
<sup>b</sup> Assumed values could not be determined from the literature and so were estimated or derived for this paper.

**Table 2:** Parameter values which varied between lambs with a description of the parameter and the reference.

| Parameter | Description | Mean | CV (dist.)<sup>a</sup> | Reference<sup>b</sup> |
| --- | --- | --- | --- | --- |
| $BW_0$ | bodyweight at birth | 3.3kg | 0.12 (n) | [1] |
| $\text{age}_{t_0}$ | age at $t_0$ | 30.8 days | 0.16 (n) | [1] |
| $b$ | Gompertz growth parameter | 0.016 per day | 0.1 (n) | Assumed |
| $f_0$ | fecundity of adult worms at $I = 0$ | 228 per day | 0.05 (g) | [2] |
| $f_1$ | fecundity of adult worms at $I = 1$ | 17 per day | 0.05 (g) | [2] |
| $C_m$ | DM feed intake for maintenance | 0.058 kg<sup>0.25</sup> per day | 0.15 (n) | [3] |
| $C_{25}$ | cum. exposure for 25% immune response | 450000 | 0.50 (g) | Assumed |
| $c_{I2}$ | DM for maintenance of immunity | 0.0035 kg<sup>0.25</sup> per day | 0.15 (n) | [4] |

<sup>a</sup> Coefficients of variation (CV) chosen to match Doeschl-Wilson et al. (2008) are shown for each parameter, where “n” denotes a normal distribution and “g” denotes a gamma distribution.
<sup>b</sup> [1] - Kenyon et al. (2013), [2] - Stear and Bishop (1999), [3] - Martins et al. (2019), [4] - Galvani et al. (2008)

**Table 3:**
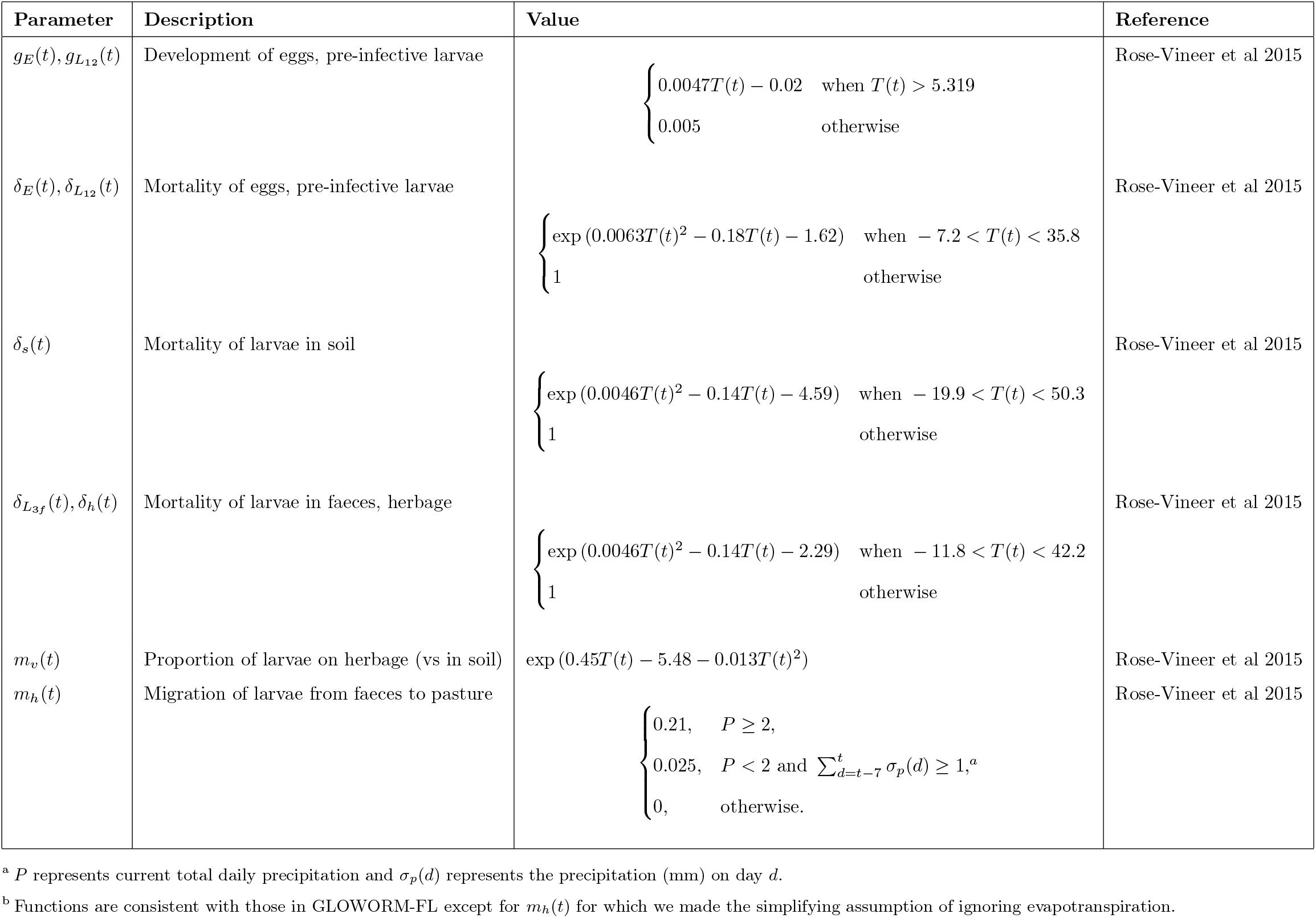
Parameter values and functional forms for the free-living parasite dynamics with a description and references.

#### 2.2.1. Body weight

We model the body weight based on the methodology proposed by Filipe et al. (2023). For ease of interpretation we first consider the body weight of a hypothetical parasite-naive animal before moving on to describe the more complex case of a parasitised animal. We assume the rate of dry matter (DM) feed intake for such a parasite-naive animal, *F*_*naive*_(*t*), expressed as a function of both body weight, *BW*_*naive*_(*t*), and rate of body weight change, 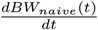, to be

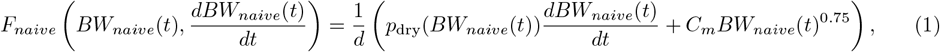

where *p*_dry_(*BW*_*naive*_(*t*)) is the proportion of growth in body weight that is DM ingested (see Eqn 3), *C*_*m*_ is the rate of biomass used for maintenance functions, *d* is the digestibility of dry matter, so that *d* · *F*_*naive*_ is the amount of DM digested per day, and the power of 0.75 on the body weight term relates to the allometric relationship that relates feed intake to body weight. We further assume that body weight growth in a parasite-naive animal follows a Gompertz function, i.e., at time *t*,

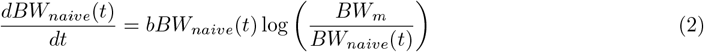

where the parameter *b* determines the rate of growth of the animal towards the mature body weight, *BW*_*m*_. Finally, we have that

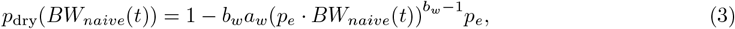

where *p*_*e*_ is the proportion of the body weight that excludes gut contents, and *a*_*w*_ and *b*_*w*_ are parameters fitted to an allometric relationship between water content and empty body weight.

The formula for the rate of DM feed intake for animal *j* in the presence of parasites, *F*_*j*_, includes additional terms describing biomass requirements for both the increase and maintenance of immunity,

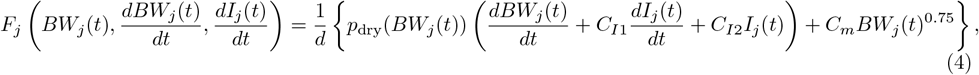

(see Sec. 2.2.5 for a description of host immunity, *I*_*j*_.) In this paper we make the same simplifying assumption as Filipe et al. (2023) to focus on the immunity-related requirements and neglect the cost of repairing worm-induced damage to the intestine, which is thought to be comparatively smaller (Houdijk et al., 2001), and we assume that dry matter requirements of the parasites are negligible. We assume that the biomass requirements for maintenance functions for a parasitised and a non-parasitised animal at the same body weight are identical. In addition, we also assume that the feed intake of a parasitised animal, *F*_*j*_(*t*), will equal the product of the feed intake of a parasite-naive animal and the anorexia coefficient, *A*_*j*_(*t*), which describes the degree to which feed intake is reduced due to parasitism, such that

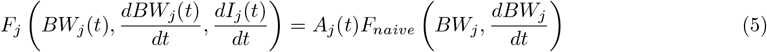

(see Sec. 2.2.6). Combining these assumptions and substituting eqns. 1 and 4 into eqn. 5 then re-arranging gives

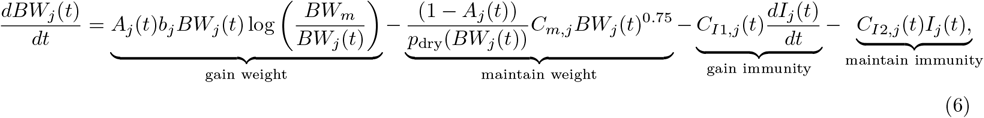

where the functions *C*_*I*1,*j*_(*t*) and *C*_*I*2,*j*_(*t*) describe animal *j*’s rates of DM biomass used to increase and maintain immunity, respectively, at time *t*, via the constants *c*_*I*1,*j*_ and *c*_*I*2,*j*_. These are

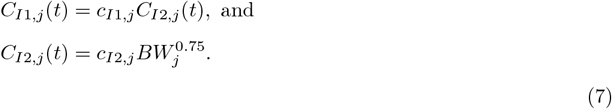

We see that in the absence of parasites 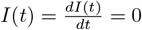 and *A*(*t*) = 1 and this reduces to the body weight gain in the parasite naive case.

#### 2.2.2. Grass growth

We assume that the grass follows the same growth pattern as in Filipe et al. (2023). Therefore, the amount of grass available for grazing at time *t, G*(*t*), with grazing taken into account is

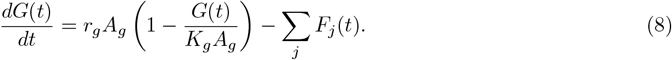

In the above, *r*_*g*_ is the growth rate of grass dry matter (DM), *A*_*g*_ is the size of the grazing area, *K*_*g*_ is the DM carry capacity of the grazing area and the summation applies over the total population of lambs. We assume that the composition and digestibility of the grass remains constant over time and thus the nutrient resources provided by the grass are assumed to remain constant. This formulation makes the assumption that grass plants are only increasing in size and not propagating in number over the growing season. We also ignore the effects of environmental conditions such as temperature and rainfall.

#### 2.2.3. Within-host parasite dynamics

The numbers of pre-adult larvae, *P*_*j*_(*t*), and adult worms, *W*_*j*_(*t*), within animal *j* at time *t*, are characterised by

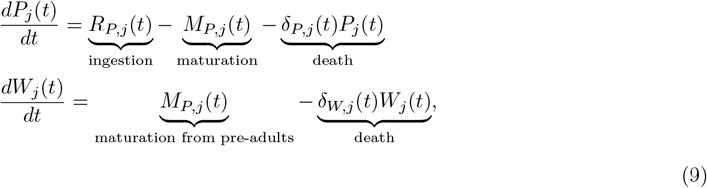

where 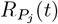 is the rate of ingestion (recruitment) of pre-adults by animal 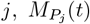 is the rate of maturation from pre-adults to adults within animal *j* (all of which are recruited into the adult stage) and δ_*P*,*j*_(*t*) and δ_*W*,*j*_(*t*) are the mortality rates of the pre-adult and adult worms, all at time *t*. The ingestion rate for animal *j* at time *t* is given by

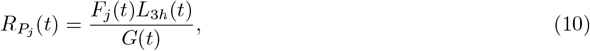

where *L*_3*h*_(*t*) is the number of infective larvae on herbage at time *t* (see Section 2.2.4). The maturation rate of pre-adult larvae at time 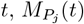, can be written as the product

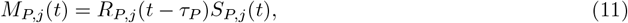

where τ_*P*_ is the duration of the pre-adult larval stage and 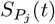 is the establishment proportion of preadult larvae in host *j*, i.e., the proportion of pre-adult larvae that survive long enough to mature into an adult worm (see Eq. 16). The pre-adult and adult worm mortality rates are each expressed as

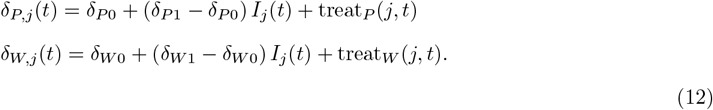

where the parameters δ_*P*0_ and δ_*P*1_ are the pre-adult mortality rates at zero and full immunity, respectively (δ_*W*0_ and δ_*W*1_ are defined equivalently). The functions treat_*P*_ (*j, t*) and treat_*W*_ (*j, t*) represent the additions to the pre-adult larvae and adult worm mortality rates due to one or several periods of anthelmintic treatment,

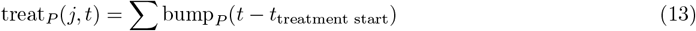

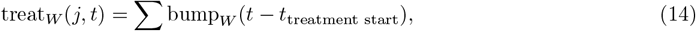

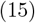

where bump_*P*_ and bump_*W*_ are functions that take the value 0 per day at *t* = 0, increase smoothly a maximum, *T*_*max*_, over a short time period (0.25 days), remain at this value for a short time day (0.5 days) and then decrease smoothly back to 0 (over a final 0.25 days). Thus treatment acts as a short pulse which clears the host of nematodes but does not provide lasting protection.

Eqs. 12 says that, in the absence of anthelmintic treatment, there is a linearly increasing relationship between the rates of pre-adult larvae or adult worm mortality within the host and the host’s current level of immunity. Finally, the establishment probability for pre-adult larvae is determined by

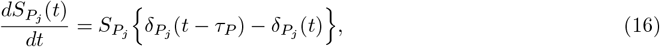

which implicitly depends on host immunity via eqns 12. This formulation of the through-stage survival comes directly from the original derivation of the time-varying DDE framework (Nisbet and Gurney, 1983).

#### 2.2.4. Free-living parasite dynamics

The state equations which correspond to the rates of change of the eggs, *E*(*t*), the grouped first and second stage larvae, *L*_12_(*t*) and infective larvae in faeces, *L*_3*f*_ can be written as

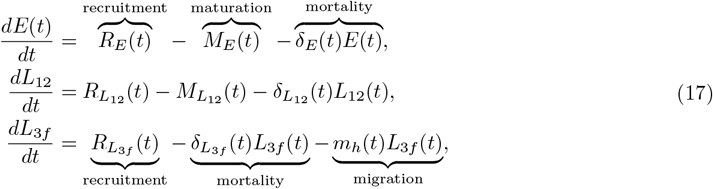

where δ_*k*_(*t*) represents the stage-specific, temperature-driven, density-independent mortality rate for stage *k* (*k* = *E, L*_12_, *L*_3*f*_), and *R*_*j*_(*t*) and *M*_*j*_(*t*) represent the number of individuals recruited into and maturing from stage *j*, respectively. Finally, *m*_*h*_(*t*) represents the rate of horizontal migration of infective larvae from faeces to pasture at time *t*. The recruitment and maturation equations can be defined by

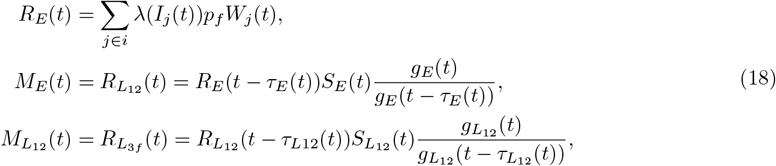

where *λ*_*j*_(*I*_*j*_(*t*)) is the fecundity of adult worms, i.e., the number of eggs produced per day by a single adult female worm within animal *j* at time *t*. This quantity is dependent on the immunity of the host, *I*_*j*_. The quantity *p*_*f*_ is the proportion of adult worms which are female and τ_*j*_(*t*), *S*_*j*_(*t*), δ_*j*_(*t*) and *g*_*j*_(*t*) are the stage duration, survival proportion, mortality rate and growth rate of individuals in stage *j* at time *t* respectively. The final fraction containing the growth rates on entry to and exit from the stage is a measure of how quickly the stage duration is changing (for full derivation see the original framework of Nisbet and Gurney (1983)). The proportion of individuals in life stage *k* that survive to the next stage, *S*_*k*_(*t*), is given by

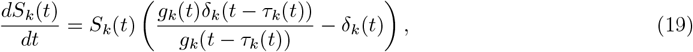

and the stage duration, τ_*k*_(*t*), is given by

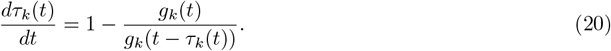

The remaining of the free-living stages is the density of larvae on pasture, *L*_3*p*_(*t*), which is split into larvae in soil, *L*_3*s*_(*t*), and larvae on herbage, *L*_3*h*_(*t*). The dynamics of larvae on pasture can be described as in GLOWORM-FL (Rose et al., 2015) such that

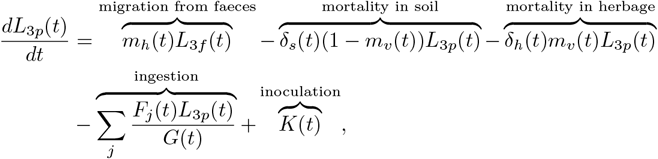

where *K*(*t*) is the rate of direct inoculation onto the patch (required to set up the initial conditions of the simulation, see Section 2.5) and *m*_*h*_(*t*) is the rate of migration of third stage larvae from faeces onto the pasture. This latter quantity depends on daily average temperature and daily amount of precipitation (for simplicity we do not consider the effect of evapotranspiration). By δ_*s*_(*t*) and δ_*h*_(*t*) we denote temperature-dependent mortality rates of third stage larvae in soil and herbage, respectively. Finally, *m*_*v*_(*t*) represents the proportion of 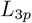 found on herbage versus soil at time *t*. This means that,

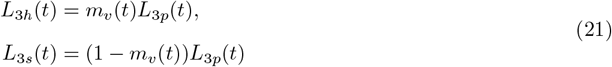

where *L*_3*h*_ and *L*_3*s*_ are the number of third stage larvae in herbage and soil, respectively. The quantity *m*_*v*_(*t*) is temperature-dependent. The rate of deposition of eggs onto pasture, *R*_*E*_(*t*), depends on the fecundity of adult worms which is influenced by the immunity, *I*_*j*_(*t*), of the host. We write

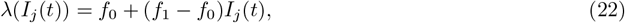

where *f*_0_ and *f*_1_ are the fecundity rates of adults worms in naive and immune hosts respectively. The parameter values for the free-living parasite dynamics are given in Table 3.

#### 2.2.5. Development of host immunity

We assume that the level of immune response in animal *j* is related to the quantity *C*_*j*_(*t*), where

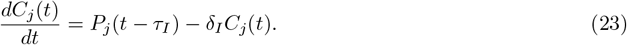

which can be described as cumulative exposure to third stage larvae in the animal’s gut, taking into account both a delay between exposure and growth of immune response of duration τ_*I*_ and the waning of the level of immunity, determined by the rate parameter δ_*I*_. It should be noted that the term “cumulative exposure” would only be completely accurate in the absence of the waning process (δ_*I*_ = 0); however as δ_*I*_ is small (δ_*I*_ = 0.002) its effects are likely to be relatively small over the first grazing season. We relate animal *j*’s cumulative exposure *C*_*j*_(*t*) to its level of immunity, *I*_*j*_(*t*) according to a von Bertalanffy-type function, which constrains the immunity level to lie between 0 and 1, as in Filipe et al. (2023), given by

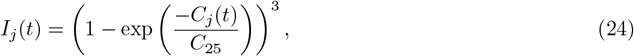

where *C*_25_ is the cumulative L3 exposure resulting in approximately 25% of the maximum immunity level being reached. The rate of change of the immunity level is

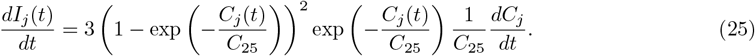

Our approach is similar to that taken in Filipe et al. (2023), in which the authors express cumulative exposure in a way that more closely resembles

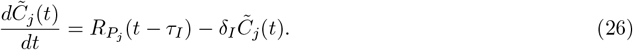

The quantity 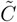 has dimension nematode, i.e. it has no time dimension, and its growth is determined by the delayed rate of ingestion 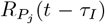, as opposed to the stage three larval burden, *P*_*j*_(*t* − τ_*I*_), as in 23. We have modified this approach since under Eq. 26 our model would allow no differential effect of targeted anthelmintic treatment on immunity of the lambs actually chosen for treatment. This is because all lambs will continue to ingest stage three larvae at a rate that is determined by their rate of feed intake and the larval density on pasture and so *C*_*j*_ (and thus acquisition and maintenance of immunity) will be unaffected by the application of the treatment in the short term. Their rate of feed intake would be affected by the reduction in the pasture larval count, due to the treated sub-population’s reduced contribution of eggs, leading to slower gain in cumulative exposure, and hence a smaller reduction in feed intake. However, this effect will be felt equally across all animals grazing on the pasture.

#### 2.2.6. Parasite-induced anorexia

In Laurenson et al. (2011) and Berk et al. (2016a), the authors describe the reduction in feed intake “RED” being proportional to the sum of the rate of change of immunity-related traits from among: establishment of larvae, worm fecundity and worm mortality. These are, in turn, related sigmoidally to the cumulative exposure to larvae. We follow the general form of this approach by assuming that the reduction in feed intake, 1 − *A*(*t*), is proportional to the rate of acquisition of immunity (which we have defined as being sigmoidally related to cumulative exposure). We further constrain *A*(*t*) to lie between 0 and 1.2, such that

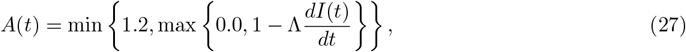

where the quantity Λ is a scaling factor.

### 2.3. Model parameterisation

We have parameterised our model based on *T. circumcincta* though the framework described could be applied to a range of parasite species by reparameterising the nematode vital rates with those appropriate for the species of interest. By parameterising the model based on empirical data and relationships from published lab and field studies we ensure that our model predictions will be generalisable to a range of farming systems and environmental conditions. We also avoid the limitations and technical challenges of fitting complex process-based models to data from a small number of field trials. The full set of parameter values and their sources are given in Tables 1 and 2. Further explanation of some of these parameters and their derivation is given below.

#### 2.3.1. Energy required for maintenance and immunity

The estimate for the energy required for maintenance, *C*_*m*_, was determined as follows. If we consider Equation 1 for a mature sheep then we can set the rate of change of the body weight to be zero and with known values for the other parameters we can calculate *C*_*m*_. Ad libitum DM intake ((Martins et al., 2019), Table 3) is 76.6 grams per day per kg^0.75^. At a mature body weight of *BW*_*m*_ = 55kg, total daily FI is 76.6 grams per day per kg^0.75^ *×* 55^0.75^ kg^0.75^ = 1.547 kg per day, and according to Eq. 1, *C*_*m*_ = 0.0766 *×* 0.756 = 0.058kg^0.25^ per day.

We determined the energy required to increase immunity, *C*_*I*1_, and maintain immunity, *C*_*I*2_, using the methodology described in Filipe et al 2022, with the allometric scaling parameters, *a*_*w*_, *b*_*w*_, *a*_*p*_ and *b*_*p*_, taken from Galvani et al 2008.

#### 2.3.2. Trait variability

The trait parameters assigned to individual animals governing their initial body weights, growth rates, DM requirements for maintenance/immunity and the fecundity of adult worms were assumed to vary between animals. This variability ensures that the parasite burden will vary between animals over the duration of the study with some animals’ body weights being more strongly affected by parasitism than others. This variability then forms the basis for selection of animals for treatment under different regimes.

The parameter values which were varied along with their means and coefficients of variation are given in Table 2. Initial body weights and times of birth were assumed to be normally distributed with means and coefficients of variation taken from the data of Kenyon et al. (2013). The coefficients of variation of the other parameter values were taken to match those assumed by Doeschl-Wilson et al. (2008).

### 2.4. Sub-model for validation against experimental challenge data

To carry out the validation against experimental challenge data (see Section 3.1) we use only part of the full model described here. Specifically, sections 2.2.2 and 2.2.4 are not required. Instead, we modify Equation 10 such that

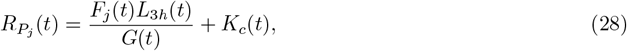

where *L*_3*h*_(*t*) = 0 because all of the free-living stages are set to be empty and *K*_*c*_(*t*) is set to take the value of *K*_0_ (= 7000 in the validation used here) on any day where a challenge is administered and 0 otherwise. To ensure that the free-living compartments of the model remain empty we can set the inoculation onto pasture, *K*(*t*), to equal 0 (Equation 21) and set the rate of deposition of eggs onto pasture *λ*(*I*_*j*_(*t*)) to equal 0. Note that we can still calculate FECs from this model by using the estimated adult worm burden, *W*_*j*_(*t*), and the estimated immunity, *I*_*j*_(*t*), extracted from the model output, as in equation 18.

### 2.5. Model history and initial conditions

The model is initialised such that all animals are considered parasite-free with

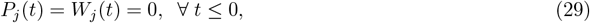

with *t* < 0 necessary due to the presence of time-delays in the model requiring that values before *t* = 0 must be referenced. Lambs are assumed to be born with no immunity such that

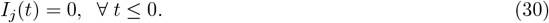

The birth weight of each animal, *j, BW*_0,*j*_ is assigned when beginning the simulation according to the values in Table 2.

We seed the pasture with *L*_0_ infective larvae at time *t* = 0 (*t*_0_) where the proportion of *L*_3*s*_ and *L*_3*h*_ being determined by the weather conditions at *t*_0_. All other free-living parasite stages are assumed to be empty at *t* ≤ 0. The grass is assumed to be at a constant level *G*_0_ at time *t*_0_ and the grass level before *t*_0_ is not required. For our simulations we assume that *L*_0_ = 2000 larvae per kg. Given the weather at this time of the year this translates to approximately 60-300 larvae on herbage, *L*_3*h*_, per kg of DM over the first 2 weeks of the study. We set *G*_0_ to be 2500kg of DM per hectare.

If we assume that the weather conditions are constant for all *t* < 0 then the stage durations for free-living stage *k*, τ_*k*0_ can be written as

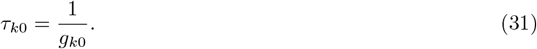

where *g*_*k*0_ is the development rate of stage *k* at *t* = 0. Similarly, the survival through the stage for stage *k, S*_*k*0_ can be written as

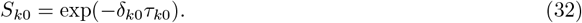

where δ_*k*0_ is the mortality rate of stage *k* at *t* = 0.

### 2.6. Husbandry system

We parameterise the body weight and feed intake components of the model based on Texel cross sheep, though the findings are generalisable to a wide variety of breeds provided the management is equivalent and the climate is similar, as only the absolute weights would change whilst the patterns of increase and decrease would remain the same. Animals are assumed to be born in early March and to start eating grass (and thus ingesting parasites) in the middle of April. We assume 60 lambs grazing at a stocking density of 24 lambs per hectare (unless explicitly stated otherwise for direct comparison with validation data). The assumed husbandry system (e.g. lambing dates, weaning date, breed of sheep) were based on the findings from the survey of Howell et al (in press).

### 2.7. Model validation

We validate the model against data from both a challenge experiment and a field experiment. We present this comparison visually and summarise the relationships between the observed and predicted body weights by calculating the repeated measures correlation coefficients. The correlation coefficients were calculated by assigning each simulated and observed lamb a ranking based on the final body weights and matching simulated body weight trajectories with observed trajectories based on these rankings. Only the body weights beginning from late May (Figures 3 and 4) were used to calculate correlation coefficients i.e. birth weights were not included. The rmcorr R package was then used to calculate correlation coefficients for each treatment group (experimental inoculation) or each combination of treatment group and paddock (field trial) accounting for correlation between repeated observations or simulations from individual lambs (Bakdash and Marusich, 2024). A full model validation would require a substantially larger dataset; however, we believe that we capture the pattern of dynamics sufficiently accurately that we explore some plausible impacts of different treatment regimes on the dynamics of the nematode population on farm.

**Figure 3:**
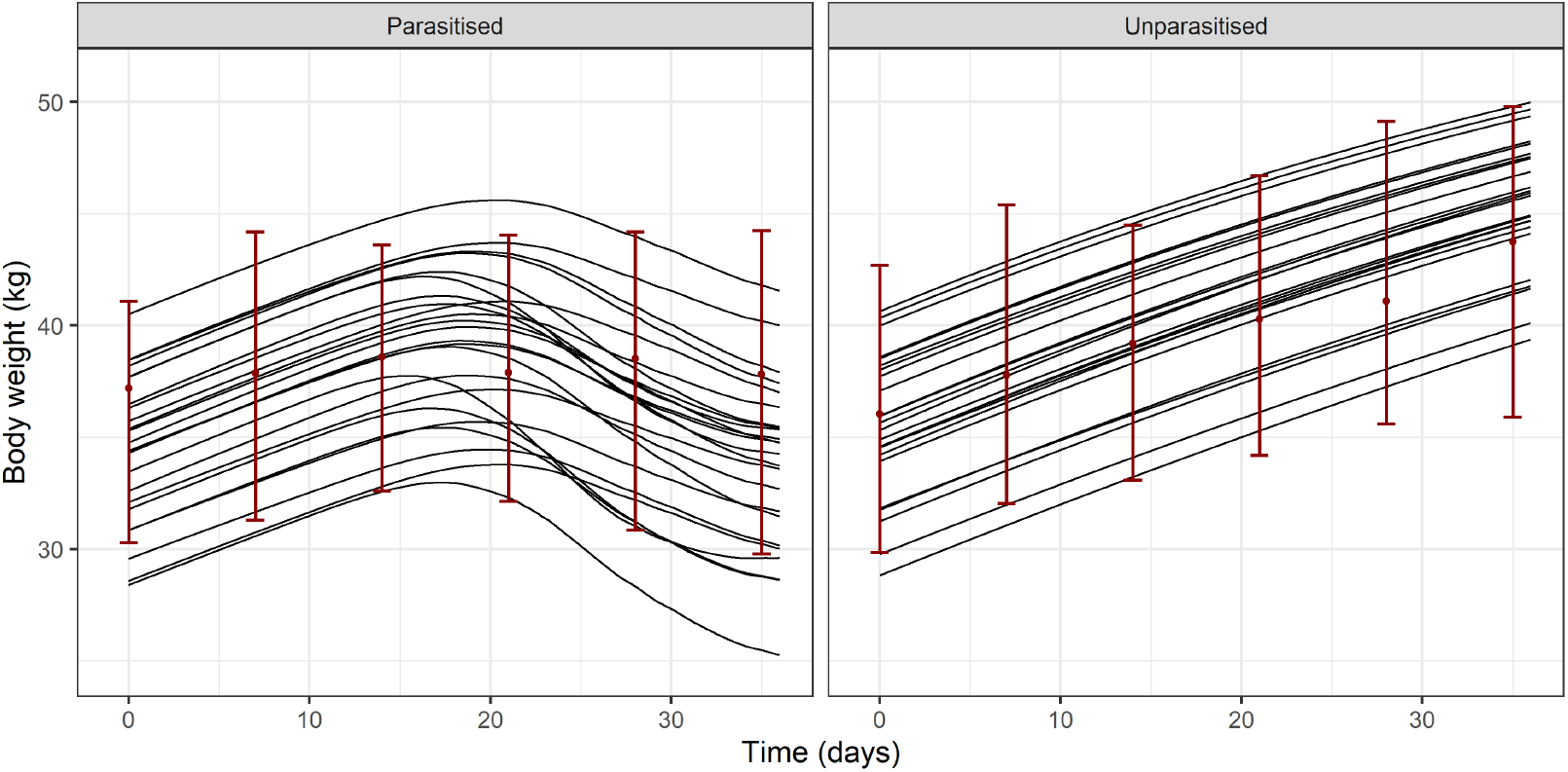
Comparison with experimental challenge data (Fox et al., 2018) based on Suffolk x mule lambs: The two panels show the model predictions of growth weights of 24 lambs (lines) that were parasitised (left) or unparasitised (right). The red points and ranges represent the median, minimum and maximum observed body weights from the challenge experiment at each observation time.

**Figure 4:**
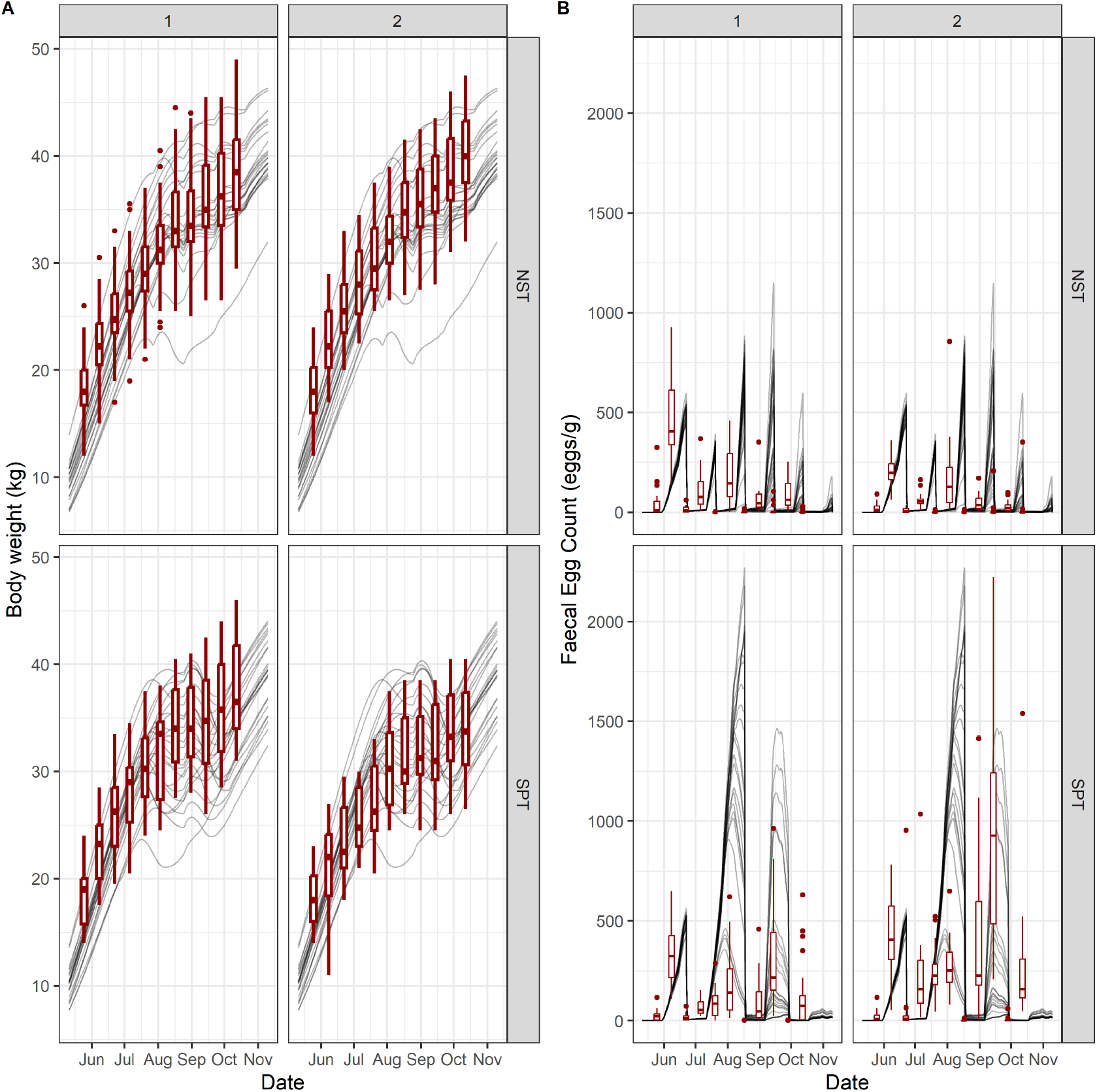
Comparison with field data (Kenyon et al., 2013): Panel A shows the model predictions of growth weights of 24 lambs (grey lines) plotted against the observed body weights of the 24 lambs (red boxplots with red points representing extreme values) on each of two paddocks (within panel) for each of the NST and SPT treatments (top and bottom). Panel B shows the model predictions of FECs for each lamb (grey lines) plotted against the observed FECs (red boxplots) on each of two paddocks (left and right) for each of the NST and SPT treatments (top and bottom).

#### 2.7.1. Comparison of predicted body weights against experimentally inoculated lambs

We compared the predicted body weights against the body weights of 48 lambs from the experiment presented by Fox et al. (2018), where 24 Suffolk x mule lambs were fed *ad lib* and challenged with 7,000 *T. circumcincta* larvae 3 times per week over a period of 35 days and a further 24 lambs control lambs were fed *ad lib* but not parasitised. In drawing these comparisons we use a simplified version of the model which ignores the free-living stages and predicts the in-host dynamics based on infective larval challenges (see Section 2.4) and so no environmental data was required here. Comparison with this challenge data allows us to isolate the in-host processes of our model for validation and to ensure that larval challenge has an appropriate effect on lamb body weights without needing to consider the numerous other processes that take place when lambs are grazing on pasture. Host modelling parameters were altered to be suitable for the animals concerned. Specifically, we increased the mature body weight, *BW*_*m*_, to 60kg (from 55kg) and increased the growth rate, *b*, to 0.018 (from 0.016) from those assumed for Texel cross due to the differences in growth rates of the breeds (Leymaster and Jenkins, 1993). We set the coefficient of variation of *C*_25_ to be 20%. *C*_25_ is very challenging to estimate but it is thought likely that the route of infection (experimental inoculation versus infection via grazing on infected pasture) may influence its value.

#### 2.7.2 Comparison of model outputs with field data

When validating against the field data (Kenyon et al., 2013) we run the model assuming 24 lambs on a one hectare patch and match timings of lambing, turnout and treatments to those in the study. We draw birth weights of lambs from a distribution with mean and coefficient of variation estimated from the birth weights observed in the field data to ensure a consistent baseline from which body weight trajectories are calculated. The field data give both body weights and FECs of cohorts of lambs under different treatment regimes at regular time intervals over a full grazing season under a UK climate, making it an ideal dataset against which to compare our model. Comparisons are made between our modelled body weights and FECs from lambs in both the neo-suppresive treatment and strategic prophylactic treatment groups (see Section 2.8 for details). We present qualitative comparisons between our model output under these treatment regimes and the field data from 2006, which is the first year of the study, because there was evidence of AR in subsequent years. We set the treatment efficacy to be approximately 99% in line with the findings of the first year of the study before anthelmintic resistance had had time to develop.

### 2.8. Anthelmintic treatment regimes

By explicitly modelling each lamb on the farm individually we are able to target anthelmintic treatment at specific lambs based on criteria, such as FECs, to reduce anthelmintic use at the flock level. We consider 4 different anthelmintic treatment regimes and the timeline of events for each regime throughout the season is shown in Figure 2. The first treatment occasion occurs after weaning in mid-June when treatment is applied to all groups (with the possible exception of group TT, as described below), after which treatments are applied to individual groups as shown (Figure 2). The four treatment regimes considered were chosen in response to a series of surveys and focus group sessions with farmers discussing viable options for anthelmintic control (Howell et al., in press) and can be described as:

- **NST:** Neo-suppresive treatment whereby all lambs are treated with anthelmintics at weaning and then every 4 weeks until they are sold to slaughter at the end of the season.
- **SPT:** Strategic prophylactic treatment whereby all lambs are treated with anthelmintics at strategically appropriate times i.e. at weaning and 6 weeks after weaning.
- **TT:** Targeted treatment where FECs are calculated for 10 lambs in the flock and treatment is applied to all animals if the mean FEC is above a specified threshold (set to 550 eggs per gram in this study).
- **T-10:** The flock is sorted in ascending order of body weight and a random 10% of animals from the top half of the flock are excluded before treating all others.

Anthelmintic treatments were assumed to induce a pulse of very high mortality lasting for one day, peaking at a value determined by the assumed (but not explicitly modelled) level of AR in the nematode population, in both pre-adult and adult worms with no lasting protection provided beyond this short pulse (see Section 2.2.3). We considered the potential efficacy of each of the proposed treatment regimes at 3 different levels of AR in the nematode population: no resistance, ~ 20% resistance and ~ 40% resistance. These resistance levels were chosen to reflect the AR shown at the end of the field trial of Kenyon et al. (2013), who observed approximately 40% resistance in the NST treatment group after 4 years and approximately 20% resistance in the SPT and targeted specific treatment (which targets treatment based on weight gain) groups under the same time frame.

### 2.9. Data accessibility

The equations were solved using an updated version of the DDE SOLVER code of Thompson and Shampine (2006) with C++ bindings such that the bulk of the model code could be written in C++. The numerical solver code and GI-NemaTracker model code are available at Benson and Ewing (2025). The validation data from the challenge and field trials could not be made available with the code but may be shared by the original authors upon request.

Our model simulations (with the exception of those simulating challenge experiments under controlled environmental conditions) use daily mean temperature and precipitation figures taken from the Copernicus Climate Data Store (Berg et al., 2021) for 2006 at the latitude 55.86 degrees N and longitude −3.20 degrees E (coinciding with the study by Kenyon et al. (2013)). This processed climate data is available in the repository with the model code.

## 3. Results

### 3.1. Comparison of predicted body weights against experimentally inoculated lambs

First we make comparisons between the body weights predicted by our model and those observed in the challenge experiment (Fox et al., 2018) (Figure 3). The pattern and range of body weights observed in both treatment groups of the challenge experiment is well captured by the model. The repeated measures correlation coefficients were estimated to be 0.92 (95% confidence interval (CI) 0.88 − 0.94) for the unparasitised lambs and 0.40 (95% CI 0.24 − 0.54) for the parasitised lambs. The lower correlation in the unparasitised group stemmed from underestimation of body weights at the later sampling points.

### 3.2. Comparison of model outputs with field data

Figure 4A compares the model predictions with the field data and shows that we capture the observed body weights well across the season both in terms of the mean and the variability between individuals, though there is evidence that we slightly underestimate them early in the year. We estimate the repeated measure correlation coefficients in the NST group to be 0.95 (95% CI 0.94 − 0.96) in paddock 1 and 0.94 (95% CI 0.93 − 0.96) in paddock 2. In the SPT group we estimate repeated measure correlation coefficients of 0.90 (95% CI 0.88 − 0.92) in paddock 1 and 0.88 (95% CI 0.84 − 0.91) in paddock 2.

In Figure 4B we capture the general pattern of the observed FECs, whereby numbers increase up to a treatment point at which time they drop to near zero before increasing again until the next treatment. In the NST treatment scenario we slightly underestimate the FECs, in large part because our predicted growth of FECs after a treatment occurs more slowly than observed in the field data. This could be because we have parameterised our model based on *T. circumcincta* but there was a range of parasite species present in the field trial. It is also possible that there is genetic variability in the nematode population that results in a distribution of pre-adult stage durations that cannot be captured by the DDE framework, which assumes a synchronous maturation to the adult stage after τ_*P*_ days. In the SPT treatment scenario we marginally underestimate the first peak in FECs, which is likely due to uncertainty surrounding the initial pasture contamination levels. We overestimate the size of the second FEC peak for many but not all animals; however, the third FEC peak is predicted with reasonable accuracy. The final peak in October is underestimated by our model, possibly due to poor understanding of environmental impacts on nematodes on pasture late in the season, as the temperature-driven migration of nematodes between soil and herbage is based on a single small study that considered a temperature range from 10-30 degrees celsius (Callinan and Westcott, 1986). It is noteworthy that our model predictions show relatively little variability between lambs early in the season, with substantially more variability after the first treatment event. This likely stems from the fact that the variation between lambs in our model is caused by intrinsic variation between lambs and their genetic traits rather than from environmental variation and stochasticity e.g. grazing different areas of a field in which the nematode population is heterogeneously distributed, which is not accounted for in the model.

Despite some disagreements between the model predictions and field observations of FECs (which are known to be highly overdispersed) we capture the body weights well and also capture the general pattern of peaks and troughs in FECs. We have not performed any fitting of our model to this field data to ensure that we do not overfit to one particular set of observations from a single study over two paddocks on a single farm in a single year.

### 3.3. Effect of treatment regimes under resistance

Figure 5 shows that in absence of resistance the NST treatment gives the highest body weights, as expected, though the T-10 regime gives very similar results due to the relatively high number of animals treated. Increasing AR results in treatments which are less effective across the different regimes, though this difference is most pronounced in the NST and T-10 groups because these regimes administer the largest number of treatments. In interpreting this result it is important to consider that resistance is expected to increase more quickly under some treatment regimes than others (i.e. resistance will be expected to increase most quickly in the NST group) so it is unlikely to be most appropriate to compare like-with-like in terms of resistance level when assessing treatment efficacy, particularly if one treatment regime is expected to lead to slower development of resistance than another. For example, it may be appropriate to compare the NST group under 40% resistance to the T-10 group under 20% resistance given the findings of Kenyon et al. (2013). This would lead us to interpret that adopting the T-10 treatment regime may result in a small loss in terms of body weight in the short term but a slight gain in the longer term due to slower emergence of resistance. However, to fully test this theory would require that our model be further extended to explicitly model the development of AR on farm over multiple seasons under each treatment regime (Berk et al., 2016b).

**Figure 5:**
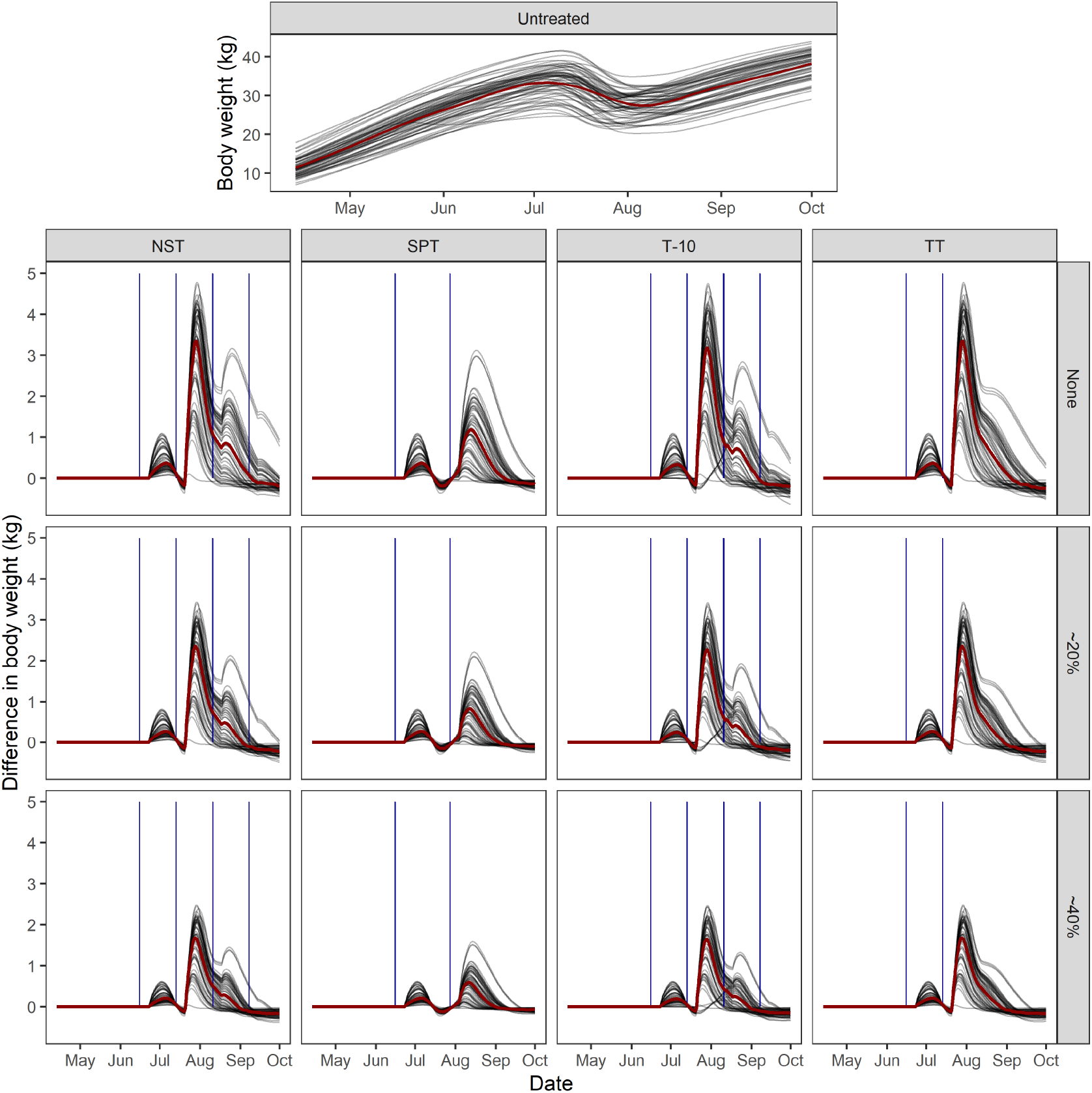
Effect of treatment regimes on body weight: The top plot shows the body weight of lambs over the grazing season without receiving any anthelmintic treatments. The lower panel of plots shows the difference in body weights between the lambs under the 4 treatment regimes (columns) and the estimated body weights of the same lambs if no treatments were applied (as shown in the top plot). The differences in body weights are assessed at the 3 specified levels of anthelmintic resistance (rows). The grey lines show the body weights (top plot) or body weight differences (bottom panel) for individual lambs and the dark red line shows a cubic regression smoothing spline (top plot) or the median body weight difference (bottom panel) across all lambs. The vertical blue lines show times at which anthelmintic treatments were administered.

Across treatments we see a general pattern that anthelmintic treatment results in a slight improvement in body weights following the first treatment in June. The difference between untreated and treated animals decreases throughout July. This is because the previously treated animals gained more body weight immediately following treatment at the expense of developing immunity courtesy of the reduced parasite burden. In July this deficit in immunity is made up for at the expense of gaining body weight at a rate slower than the untreated animals with stronger immunity. Throughout August the treated animals then show a second peak in their difference in body weight because they are challenged with fewer nematodes than those in the untreated group because previous anthelmintic application has resulted in a reduced parasite burden on pasture. The differences in body weights between predictions for the simulated untreated and treated lambs were estimated to be approximately zero by the end of September.

It should be noted that the NST and TT approaches result in equivalent management practices until the third treatment time in this case, after which point the two approaches diverge in terms of the treatments offered and predicted weight gains. Comparison of these two approaches therefore makes it clear that the treatments in September and potentially also in August may be somewhat redundant, due to high levels of immunity in the lambs and the decline in the nematode population at the end of the season resulting from worsening environmental conditions.

The FECs of each animal through time are shown in Figure 6 and the pattern of seasonal abundance of nematodes on pasture is shown in Figure 7. The differences in pasture contamination across the treatment groups and resistance scenarios match what would be expected given the impacts of the treatments on body weights, with relatively lower pasture contamination generally coinciding with higher body weights at a given point in the year; however, the differences in pasture contamination across treatment groups are less pronounced than those in body weight because pasture contamination is strongly influenced by current and previous environmental conditions (Morgan and van Dijk, 2012). We predict a relatively late peak in *L*_3*h*_, though our predictions are consistent with the GLOWORM-FL model (Rose et al., 2015). Any potential overestimation late in the year may be due to poor understanding of development and mortality rates of free-living stages of *T. circumcinta* at lower temperatures, with functions parameterised based on a small number of studies that did not consider temperatures between 4 and 16 degrees celsius which contains the bulk of the environmental conditions experienced at our study site (Pandey et al., 1989, 1993). The FECs drop in all groups at the end of the season due to the high levels of immunity in the lambs.

**Figure 6:**
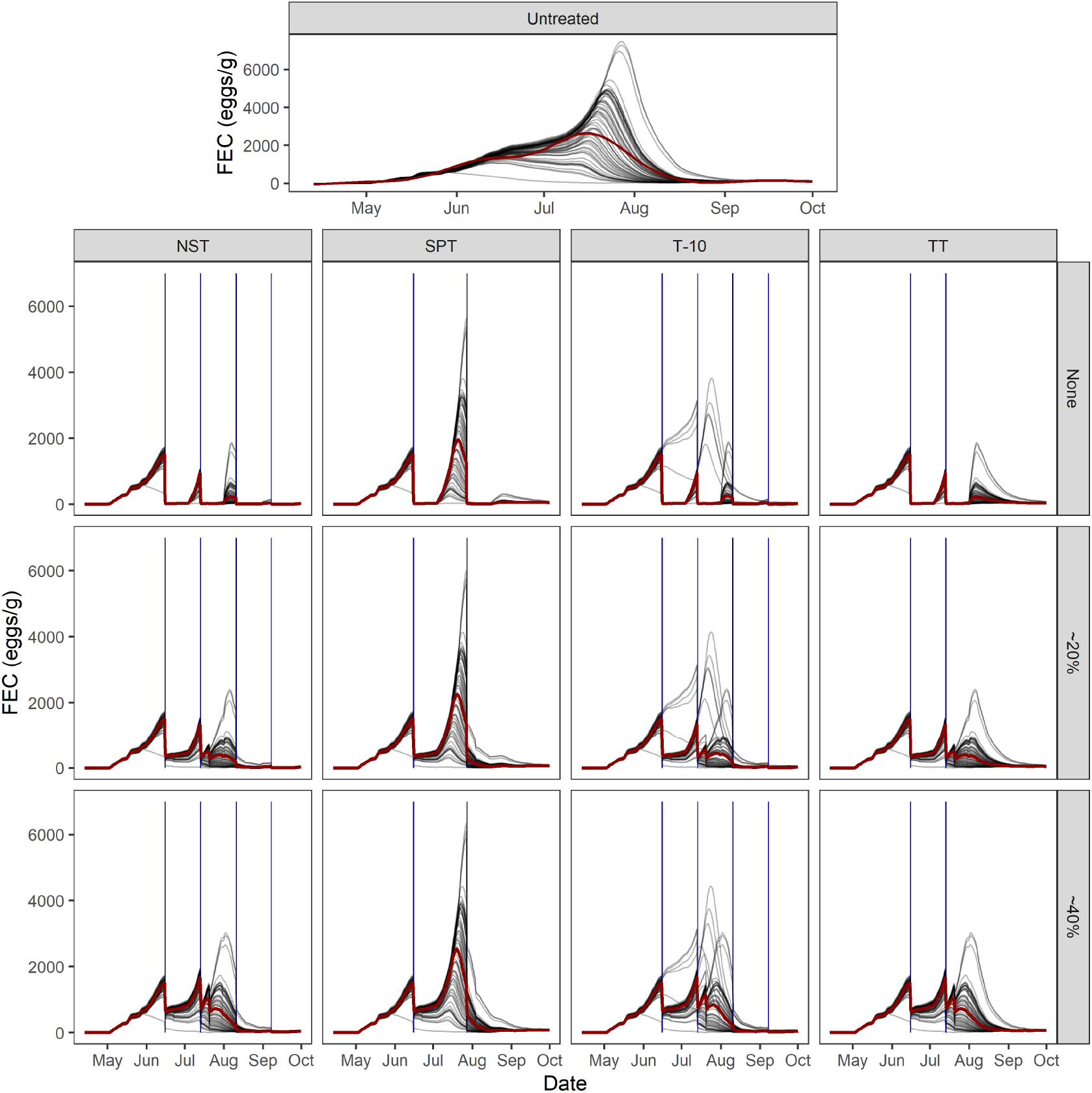
Effect of treatment regimes on FECs: The plots show the FECs of all animals through time under the 4 treatment regimes and for untreated animals (columns) at the 3 specified levels of AR (rows). The vertical blue lines show the timing of application of anthelmintic treatments.

**Figure 7:**
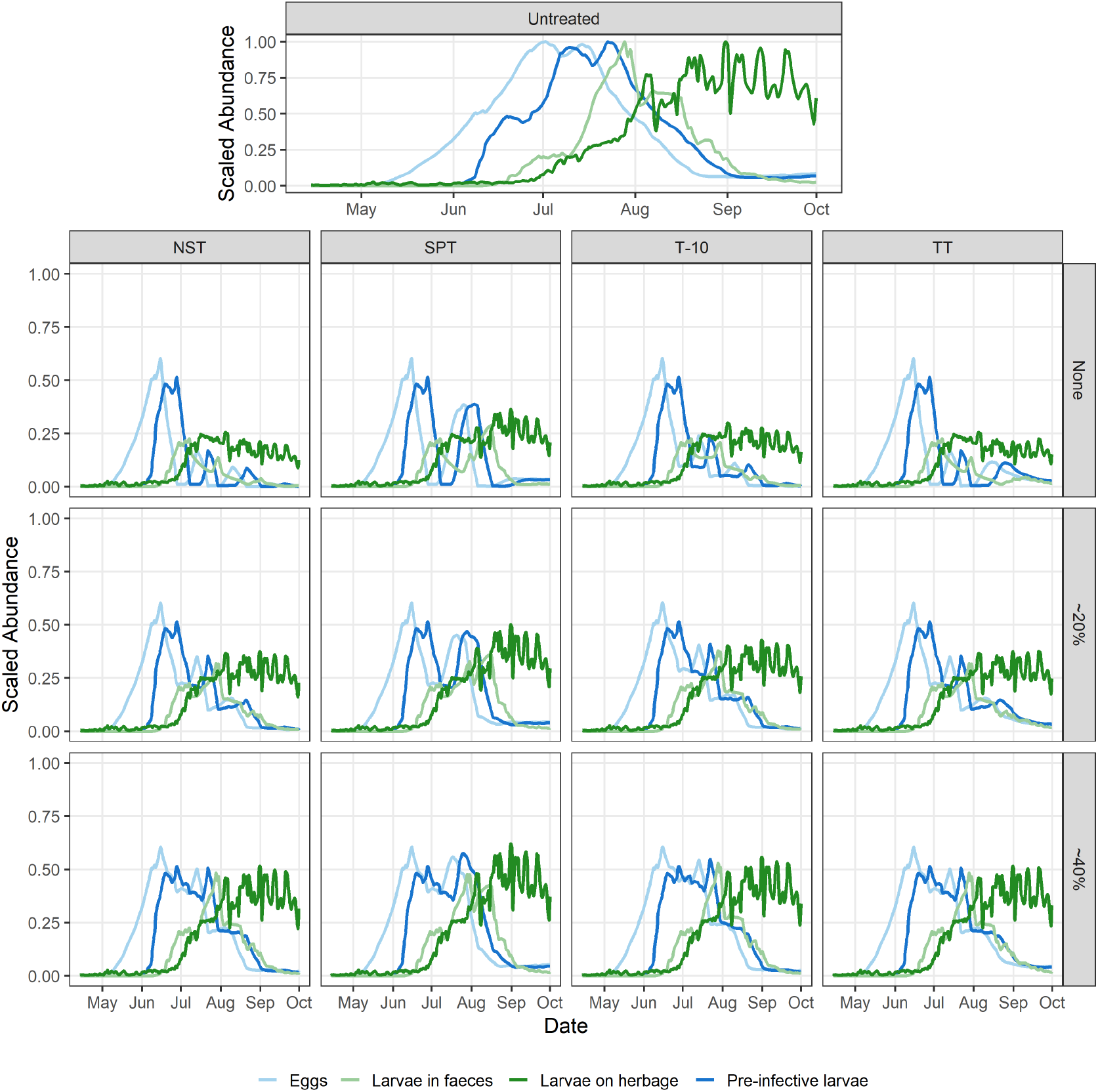
Effect of treatment regimes on pasture contamination: The plots show the abundance of eggs, pre-infective larvae (*L*_12_), larvae in faeces (*L*_3*f*_) and infective larvae on herbage (*L*_3*h*_) scaled by the maximum value of each of these life stages under the no treatment scenario. Plots are shown across the 4 treatment regimes (columns) at the 3 specified levels of AR (rows). Scaled abundances are plotted to show the relative timings of the peaks in abundance of each life stage due to the large difference in scales of the raw abundances across the life stages.

## 4. Discussion

We develop a systems modeling framework that integrates decades of research into a tool that enables in silico assessment of the relative impacts of current and potential farm system management protocols on lamb body weight gain and parasite contamination levels on farm. The mathematical model presented captures parasite dynamics both in host and on pasture and predicts the parasite burdens and consequent body weights of a phenotypically diverse population of lambs over their first grazing season. Our model results have been validated both against data from a challenge experiment, where lambs were directly inoculated with infectious larvae (Fox et al., 2018), and against data from a field study (Kenyon et al., 2013) and successfully recreates observed body weights and nematode population dynamics (Figures 3 & 4). This validated model framework was used to explore the potential efficacy of 4 anthelmintic treatment regimes that were co-created through a series of workshops and farmer focus groups. We showed that a targeted approach whereby 10% of animals are left untreated at each treatment occasion can result in only a small difference in body weights when compared to an NST approach in which all animals are treated (Figure 5). We also show that treatments administered late in the season when the lambs have relatively high immunity and conditions are less favourable for the nematodes may be relatively ineffective. If these more targeted approaches were to reduce the development of AR on farm, then this could lead to higher average weights in future years, dependent on the degree to which development of AR was slowed. By adopting a farm system level approach we have shown that we can extend on previous modelling works that have studied these processes in isolation (Vagenas et al., 2007b; Laurenson et al., 2013; Rose et al., 2015; Rose Vineer et al., 2020; Filipe et al., 2023) to predict the impact of novel control strategies that directly target individual animals whilst considering the entire environmentally-driven parasite life cycle.

We have focused on targeted treatment based on FECs as the novel control approach by which GINs could be controlled whilst potentially slowing development of AR; however, the model and codebase have been designed to be sufficiently flexible to allow ready extension to study alternative control measures. We explicitly model the weight gain of animals through time, allowing TST approaches such as the Happy Factor™algorithm, which uses individual animal weight predictions to determine animals requiring treatment (McBean et al., 2021), to be straightforwardly included in future model comparisons. In this paper we have focused on a set stocking scenario but the explicit inclusion of weather-driven time delays in development of nematodes on pasture via the DDE model makes our approach ideally suited to exploring alterative approaches to controlling nematode infections. For example, mob grazing is thought to reduce nematode burdens in sheep but the timing of return to a previously grazed patch is likely to strongly influence efficacy (Louie et al., 2006; Colvin et al., 2012; Walkden-Brown et al., 2013). Our model code has been written to facilitate future extensions that explicitly allow movement of animals between paddocks at predefined times such that we could test potential efficacy of mob grazing approaches.

It has been hypothesised that climate change, which is predicted to result in wetter, milder winters and hotter, drier summers in the UK, will affect the seasonal abundance of parasitic nematodes in the future with evidence of some such changes having already emerged (van Dijk et al., 2010; Fox et al., 2015). We have not focused on exploring the likely impacts of intra- and inter-annual variation in environmental conditions in this paper, though these factors may affect the optimum timing of treatments due to their impacts on the parasite burden on pasture. Nonetheless, by explicitly modelling the interaction between environmentally-forced free-living parasites and individual host animals our model and straightforward extensions of it can be used to address three of the research priorities to understand climate change challenges to farmed ruminants in temperate regions identified by van Dijk et al. (2010), namely: quantification of the overall effects of climatic changes on parasite dynamics, identification of likely adaptation of farm management to climate change and continuing research into alternative strategies for parasite control.

At present we model the effects of resistance by adjusting the efficacy of anthelmintic treatments to give the desired level of mortality under each resistance scenario. To fully understand which novel control scenarios may give the best balance between controlling nematode populations and slowing the development of AR would require that we also model the emergence of resistance over time. Previous models have predicted that targeted treatment approaches can improve average daily weight gain in calves whilst limiting the emergence of AR (Berk et al., 2016b). Model extensions could draw from previous work modelling the emergence of AR in livestock (Smith, 1990; Gaba et al., 2006; Berk et al., 2016b) as well as the extensive body of literature regarding modelling the emergence of antimicrobial resistance (Niewiadomska et al., 2019).

Our model estimates that body weights at the end of September will be approximately equal across the groups and between treated and untreated animals. This is consistent with the findings of Kenyon et al. (2013), who showed no statistically significant difference in body weights 22 weeks after turnout in NST, SPT and TST treatment groups in any year over a 5 year field trial. In considering that the eventual body weights are equivalent across the treatment groups it is important to note that the model does not consider the secondary negative health consequences (e.g. clinical gastro-enteritis) that could be associated with body weights being up to 5kg lighter in late July and early August in an untreated scenario than in some of the treatment scenarios presented here. Such secondary health implications may mean that all animals do not recover to the body weight we predict due to unmodelled processes. We also only consider losses in body weight gain due to the nutrient resources required to gain immunity and do not explicitly model a damage and repair mechanism stemming from parasitism, which may result in larger differences late in the season, though the energy lost to repairing damage is though to be substantially less than that required to mount an immune response (see Section 2.2 for a discussion of this assumption).

We have developed a farm system level mathematical model that links together the in-host and free-living stages of the GIN life cycle whilst accounting for the impacts of climatic drivers on the free-living stages and the relationships between worm burden and immunity of a population of lambs including trait diversity. We show that FEC-based TT approaches can result in similar performance at a flock level to more intensive treatment regimes, lending support to studies which have proposed such approaches as desirable control options (Kenyon and Jackson, 2012; Charlier et al., 2014). The model can be readily employed with minimal modification to answer further pertinent questions such as how climate change might affect the burden of GINs on sheep in the future or how a broader range of sustainable worm control practices could be implemented to control nematode infestations under different environmental conditions (van Dijk et al., 2010; Charlier et al., 2018). The individual-based modelling framework also gives the flexibility to explore the effectiveness of different sustainable worm control approaches in a range of different farming systems, taking us closer to developing bespoke models that capture differences between farms to facilitate effective knowledge exchange (Jack et al., 2017).

